# ARC-Impute: robust reference-guided spatial gene imputation through regularised cell-to-spot mapping

**DOI:** 10.64898/2026.11.17.767312

**Authors:** Ethan Han, Andrew Zhang, Jinman Kim, Jean Yang

## Abstract

**Motivation:** Reference-guided spatial imputation uses single-cell RNA sequencing references to predict genes that are unmeasured in spatial transcriptomics data. Limited shared-gene overlap can leave the mapping of reference cells to spatial locations ambiguous: different assignments may reconstruct observed genes similarly yet yield different imputed expression and cell-type estimates. This gives rise to local assignment ambiguity, source-target mismatch and reference-transfer instability.

**Results:** We present ARC-Impute, which regularises cell-to-spot mapping through anchor-residual optimisation, target-guided reference curation and routing, and twin-panel consistency. Across 17 human and mouse datasets spanning spot-based and single-cell technologies, ARC-Impute ranked first for held-out Pearson correlation throughout and exceeded the dataset-specific baseline mean by 47.3% on average. It also achieved the strongest spot-level cell-type composition recovery across three pseudospot benchmarks and retained leading accuracy and consistency under reference and training-gene perturbations. Cross-validated predictions revealed systematic differences in gene imputability.

**Availability and implementation:** Code and workflows at https://github.com/ethanhaan/arc-impute.

## 1. Introduction

Spatial transcriptomics (ST) measures gene expression within intact tissue, preserving spatial coordinates and revealing how molecular signals vary across anatomical regions and cellular neighbourhoods. However, many ST assays capture only part of the transcriptome due to either targeted gene panels or sparse transcript capture (Biancalani et al. 2021). In spot-based platforms, each spot often captures a mixture of several cells rather than a true single-cell profile (Miller et al. 2022). Single-cell RNA sequencing (scRNA-seq) offers a complementary view: it provides transcriptome-wide information at single-cell resolution, but loses spatial context during tissue dissociation (Macosko et al. 2015). Together, ST and scRNA-seq provide complementary views of tissue biology, motivating the use of scRNA-seq as a reference to predict genes that were unmeasured or were poorly captured within spatial data. In this study, we refer to this task as reference-guided spatial imputation.

Existing methods for reference-guided spatial gene imputation generally learn the relationship between scRNA-seq and ST in one of two ways: by building a shared representation or predictive model across modalities (Abdelaal et al. 2020, Chen et al. 2021), or by learning an explicit mapping from reference cells to spatial locations (Cang and Nie 2020, Biancalani et al. 2021, Vahid et al. 2023). In both cases, the relationship is learned from genes measured in both the scRNA-seq reference and the spatial dataset. Because this overlap is often limited, the correspondence between the scRNA-seq reference and spatial data is weakly constrained (Li et al. 2022). For methods that explicitly map reference cells to spatial locations, multiple mappings can reconstruct the measured genes similarly well, while producing different predictions for unmeasured genes and different estimates of spot-level cell-type composition. This weakly constrained mapping can therefore lead to incorrect reference-cell assignments and inaccurate gene expression imputation despite good reconstruction of the measured genes.

This gives rise to three distinct challenges. (i) Local assignment ambiguity arises because the informativeness of the observed genes varies across the tissue: while some spots clearly support a plausible reference mixture, sparse, noisy or spatially incoherent spots allow a flexible mapping to overfit weak local evidence, leading to inaccurate held-out predictions. (ii) Source-target mismatch arises when the scRNA-seq reference does not match the spatial tissue. If some cell types or states are over- or under-represented in the reference, the model may assign cells because they are available in the reference rather than because they are supported by the spatial data. (iii) Reference-transfer instability arises because the mapping is sensitive to the shared genes and reference cells used during fitting. Because these provide only a partial view of the cross-modal relationship, reasonable variations to either may produce different assignments and thus different imputed gene expression and cell-type estimates. Existing methods address parts of this challenge by improving cross-modal alignment or by constraining the inferred mapping through spatial density, transport cost or cell-type abundance (Abdelaal et al. 2020, Cang and Nie 2020, Biancalani et al. 2021, Chen et al. 2021, Vahid et al. 2023, Li et al. 2025). However, no existing approach jointly addresses all three sources of instability: local assignment ambiguity, source-target mismatch and reference-transfer instability.

Here, we introduce ARC-Impute, a reference-guided spatial imputation model designed to address these three problems. Building on Tangram’s probabilistic cell-to-spot mapping formulation (Biancalani et al. 2021), ARC-Impute learns the cell-to-spot assignment through three complementary modules that jointly structure and regularise the mapping: (i) anchor-residual optimisation, which stabilises the assignment around a coarse cell-type-to-region anchor while allowing local refinement; (ii) reference curation and assignment routing, which reduce bias from reference populations that are poorly supported by the spatial data; and (iii) twin-panel consistency, which reduces sensitivity to shared-gene choice by encouraging mappings learned from alternative gene panels to agree. The resulting mapping is used to predict unmeasured spatial gene expression and estimate spot-level cell-type composition. We evaluate ARC-Impute across 17 datasets using held-out gene reconstruction, pseudospot deconvolution and robustness analyses under reference resampling and gene-panel perturbation. We further use cross-validated predictions across reference-guided models to identify genes that are consistently easier or more difficult to impute and determine whether these differences reflect specific biological programmes.

## 2. Materials and methods

### 2.1. ARC-Impute formulation

Throughout this formulation, *spot* refers broadly to either a capture spot or a single-cell spatial observation.

ARC-Impute jointly predicts unmeasured spatial gene expression and estimates spot-level cell-type composition by learning a structured probabilistic mapping from an annotated scRNA-seq reference *X* to a spatial target *Y*. Because the shared genes only partially determine the cell-to-spot correspondence, ARC-Impute constrains the mapping against local assignment ambiguity, source-target mismatch and reference-transfer instability rather than allowing reconstruction fit alone to determine the assignment. As illustrated in Figure 1, target-derived cell-type demand first curates the reference to obtain 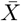 and defines the routing bias *b*; a coarse biological anchor *A* provides initial spatial support; and two overlapping gene panels *G*_1_ and *G*_2_ produce branch-specific mappings *M*_1_ and *M*_2_.

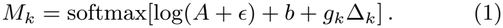

**Figure 1.**
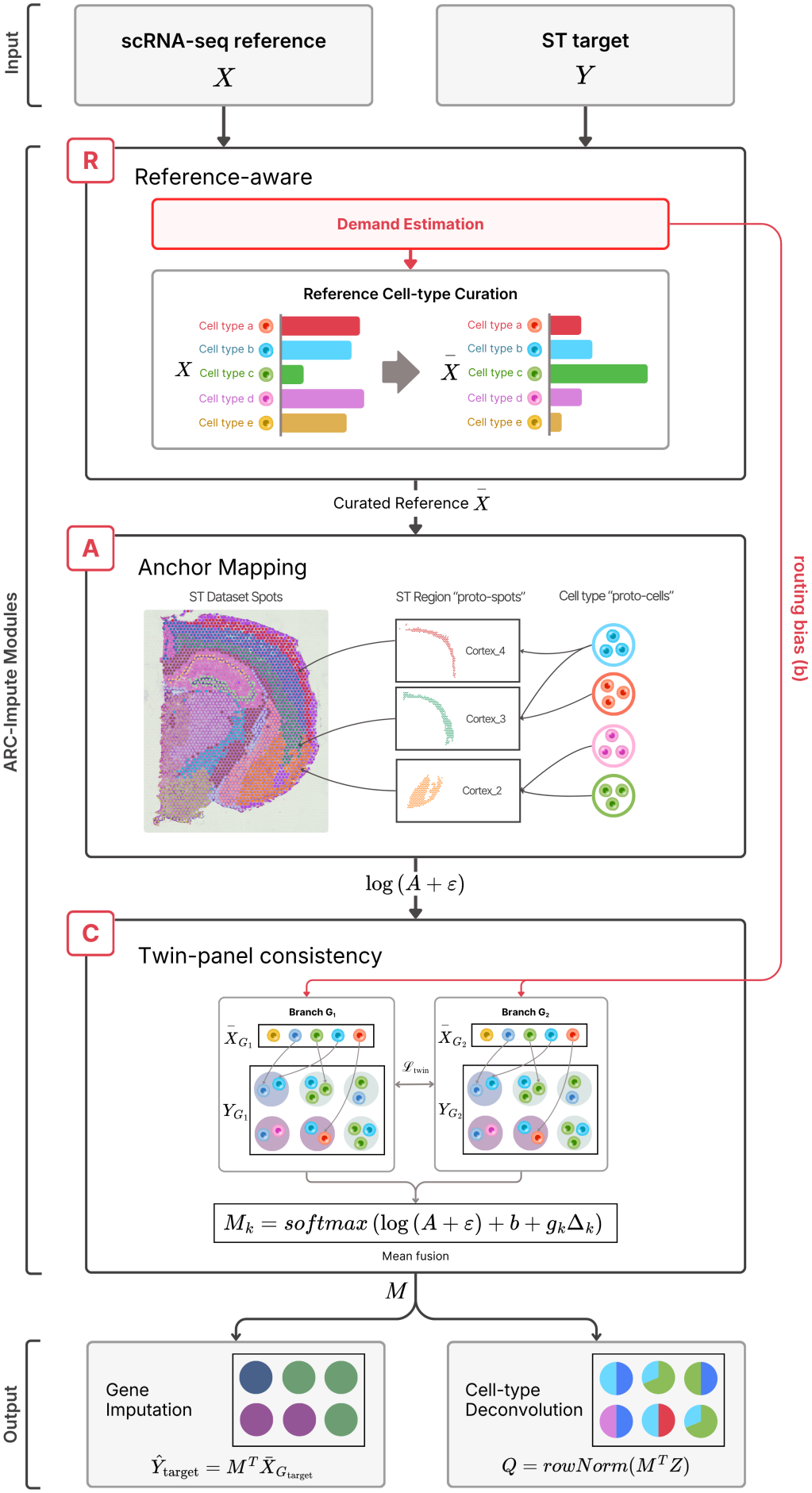
Overview of ARC-Impute. Target-derived demand guides reference curation and assignment routing (R), a coarse biological anchor constrains the cell-to-spot mapping (A), and twin-panel consistency regularises mappings learned from complementary gene panels (C). The fused mapping supports spatial gene imputation and spot-level cell-type deconvolution.

Here, *M*_*k*_ ∈ ℝ^*N×S*^ is the branch-specific mapping for *k* ∈ {1, 2}, and *A* ∈ ℝ^*N×S*^ is the coarse cell-to-spot anchor mapping, where *N* is the number of curated reference cells and *S* is the number of spatial spots. *b* ∈ ℝ^*N×S*^ is the fixed routing bias, Δ_*k*_ ∈ ℝ^*N×S*^ is the learned residual for branch *k*, and *g*_*k*_ is a learned scalar controlling its size. *ϵ* is a small constant for numerical stability. The softmax is applied across spatial spots for each reference cell, so its mapping weights sum to 1. The anchor *A* provides a coarse assignment of reference cells to spatial spots, while *b* adjusts this assignment using target-supported cell-type demand. The learned residual *g*_*k*_Δ_*k*_ then allows each branch to refine the mapping without moving too far from the anchor. The two branches share the same anchor and routing structure but learn separate residuals from *G*_1_ and *G*_2_, allowing agreement between the two gene panels to regularise the final mapping.

Each branch reconstructs its corresponding observed spatial expression as 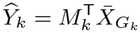 Here, 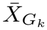 is the curated reference restricted to branch gene panel *G*_*k*_, while 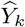 and 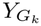 are the predicted and observed spatial expression matrices, respectively.The two mappings are jointly optimised using:

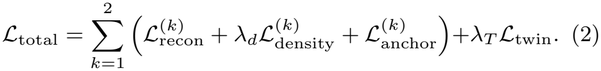

The reconstruction and density components use the same general forms as Tangram (Biancalani et al. 2021): 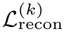is the negative mean gene-wise cosine similarity, while 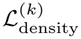 is the KL divergence between observed and predicted spot-density distributions. ARC-Impute introduces 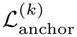 which limits residual updates and deviations from the anchor, and *L*_twin_, which enforces agreement between the two branch-specificcell-type compositions. Across experiments, the density, gate, anchor-divergence and twin-consistency regularisation weights were *λ*_*d*_ = 1, *λ*_*g*_ = 0.01, *λ*_KL_ = 0.03, and *λ*_*T*_ = 0.05, respectively, with unit weight on reconstruction; all were fixed across datasets.

After optimisation, the branch assignments are fused into a consensus mapping *M* = (*M*_1_ + *M*_2_)*/*2, which is used to transfer reference-only gene expression and aggregate reference-cell annotations,

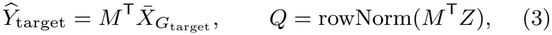

where *G*_target_ is the set of reference-only genes to be imputed, rowNorm normalises each row to sum to one, *Z* contains one-hot reference cell-type labels, and *Q* is the resulting spot-level cell-type composition.

### 2.2. ARC-Impute modules

#### Anchor-residual optimisation

To reduce overfitting where shared-gene evidence is weak, ARC-Impute learns each cell-to-spot mapping around a coarse biological anchor. Curated reference cells are averaged within annotated cell types to form pseudo-cells, and target spots are averaged within spatial regions to form pseudo-spots. A coarse cell-type-to-region mapping *P* ∈ ℝ^*C×R*^ is fitted between these aggregated profiles, where *C* and *R* are the numbers of cell types and spatial regions, respectively, and *P*_*c,r*_ represents the assignment mass from cell type *c* to region *r*. This coarse mapping is expanded uniformly within each region to form the cell-to-spot anchor *A* ∈ ℝ^*N×S*^,

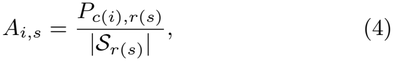

where *N* and *S* are the numbers of reference cells and spatial spots, *c*(*i*) is the type of reference cell *i, r*(*s*) is the region containing spot *s*, and *S*_*r*_ = *{t* : *r*(*t*) = *r}* is the set of spots in region *r*. Thus, each spot within a region receives an equal share of the corresponding regional assignment mass. To reduce sensitivity to initialisation, independently fitted anchors are median-aggregated element-wise and subsequently row-normalised. During branch optimisation, *A* remains fixed and each branch learns only a gated residual *g*_*k*_Δ_*k*_, with deviation controlled by

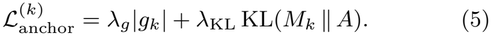

The anchor preserves stable regional structure, while the residual permits finer spot-level refinement when supported by the data.

#### Reference curation and assignment routing

To reduce source-target mismatch, ARC-Impute estimates which cell types are supported by the spatial target, then uses this information to resample the reference and guide cell-to-spot assignment. Let *µ*_*c*_ denote the shared-gene centroid of reference cell type *c*. For each spot *s*, centroid similarity is converted into a local cell-type demand distribution,

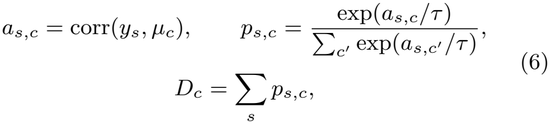

Here, *y*_*s*_ is the shared-gene expression vector for spot *s*, and corr denotes Pearson correlation. The parameter *τ* controls the sharpness of the local demand, and *D*_*c*_ aggregates support for cell type *c* across the tissue. Global demand is normalised and blended with the original reference composition *s*_*c*_ to avoid prematurely removing weakly detected populations,

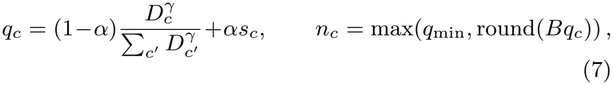

where *q*_*c*_ is the cell-type target proportion used to resample the reference, *α* ∈ [0, 1] controls the contribution of the original reference composition, *γ >* 0 controls demand sharpening, *B* is the reference-cell budget, *q*_min_ is the minimum number of retained cells per type, and *n*_*c*_ is the resulting number retained in the curated reference 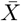. This reduces the influence of overrepresented or unsupported source populations while preserving a minimum representation of each cell type. The local demand also defines the routing bias *b*_*i,s*_ = *λ*_*R*_*p*_*s,c*(*i*)_, where *λ*_*R*_ controls the strength of this bias. This gives reference cells higher assignment weights at spots whose expression is compatible with their annotated type, while allowing residual optimisation (*g*_*k*_Δ_*k*_) to determine their final placement.

#### Twin-panel consistency

To reduce reference-transfer instability under changes to the shared-gene panel and improve the robustness of the inferred cell-to-spot mapping, ARC-Impute divides the shared genes into two overlapping panels containing a common informative core and distinct remaining genes, *G*_1_ =*G*_core_ *∪ E*_1_ and *G*_2_ = *G*_core_ *∪ E*_2_. Genes are ranked by variation across reference cell-type centroids, with the most informative retained in *G*_core_ as a shared basis and the remaining genes divided between *E*_1_ and *E*_2_. The two branches share the anchor *A* and routing bias *b*, but learn independent gated residuals from their respective panels. Agreement between the resulting mappings is enforced at the spot-level cell-type composition level through ℒ_twin_= MSE(*Q*_1_, *Q*_2_), where 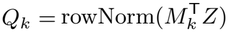 and *Z* contains one-hot reference cell-type labels. This permits reference cells of the same annotated type to be assigned interchangeably. Following optimisation, the branch mappings are mean-fused as *M* = (*M*_1_ + *M*_2_)*/*2 to obtain the final assignment used for gene imputation and cell-type deconvolution.

### 2.3. Datasets and benchmark design

We benchmarked ARC-Impute across 17 spatial transcriptomics datasets spanning human and mouse tissues, healthy and diseased states, and diverse anatomical contexts. These comprised ten spot-based datasets, including three pseudospot benchmarks constructed from native cell-resolved spatial transcriptomics data, and seven native cell-resolved datasets. Each spatial target was paired with an appropriate scRNA-seq reference, and ARC-Impute was compared against novoSpaRc, Tangram, SpaGE, SpaIM, stImpute, CytoSPACE, Seurat and LIGER. For most datasets, shared marker genes were identified by taking the deduplicated union of the top 100 genes ranked for each annotated reference group; where the overlapping gene set was limited, all available shared genes were retained. The resulting gene panel was partitioned into five folds for held-out reconstruction, with PCC used as the primary metric and cosine similarity and SSIM as complementary measures. The three pseudospot datasets additionally enabled evaluation against known spot-level cell-type proportions, while a 14-dataset perturbation benchmark assessed robustness to training-gene subsampling and reference-cell resampling. Full dataset accessions, preprocessing, reference pairings, pseudospot construction and metric definitions are provided in the Supplementary Methods.

## 3. Results

### 3.1. ARC-Impute improves spatial gene imputation

On the Human Breast Cancer Visium dataset, we benchmarked ARC-Impute against eight methods using five-fold cross-validation over shared marker genes. Held-out measured genes were predicted from the remaining genes and scRNA-seq reference, then evaluated using mean per-gene PCC, with cosine similarity and SSIM as complementary metrics (see Supplementary Methods, Section S2.1). ARC-Impute achieved the highest PCC of 0.272, compared with 0.235 for SpaIM (Figure 2a), an absolute improvement of 0.038 and relative gain of 16.0%. At the individual-gene level, ARC-Impute produced an overall upward shift in held-out-gene PCC relative to the baseline methods (Figure 2b). This indicates that the improvement extended broadly across the evaluated gene panel rather than being driven by a small number of genes. For representative held-out genes, ARC-Impute recovered major tissue-associated expression patterns while avoiding the fragmented or over-smoothed predictions produced by several baselines (Figure 2c). These genes included DST, COL6A2, MMP2 and PECAM1, linked respectively to cytoskeletal and YAP regulation, collagen-VI-associated signalling, extracellular-matrix remodelling and tumour vasculature (Jain et al. 2019, Scordamaglia et al. 2025, Ueno et al. 1997, Horak et al. 1992). Across these examples, ARC-Impute more closely reproduced the observed spatial expression patterns across the tissue, consistent with the quantitative improvements in gene-level imputation.

**Figure 2.**
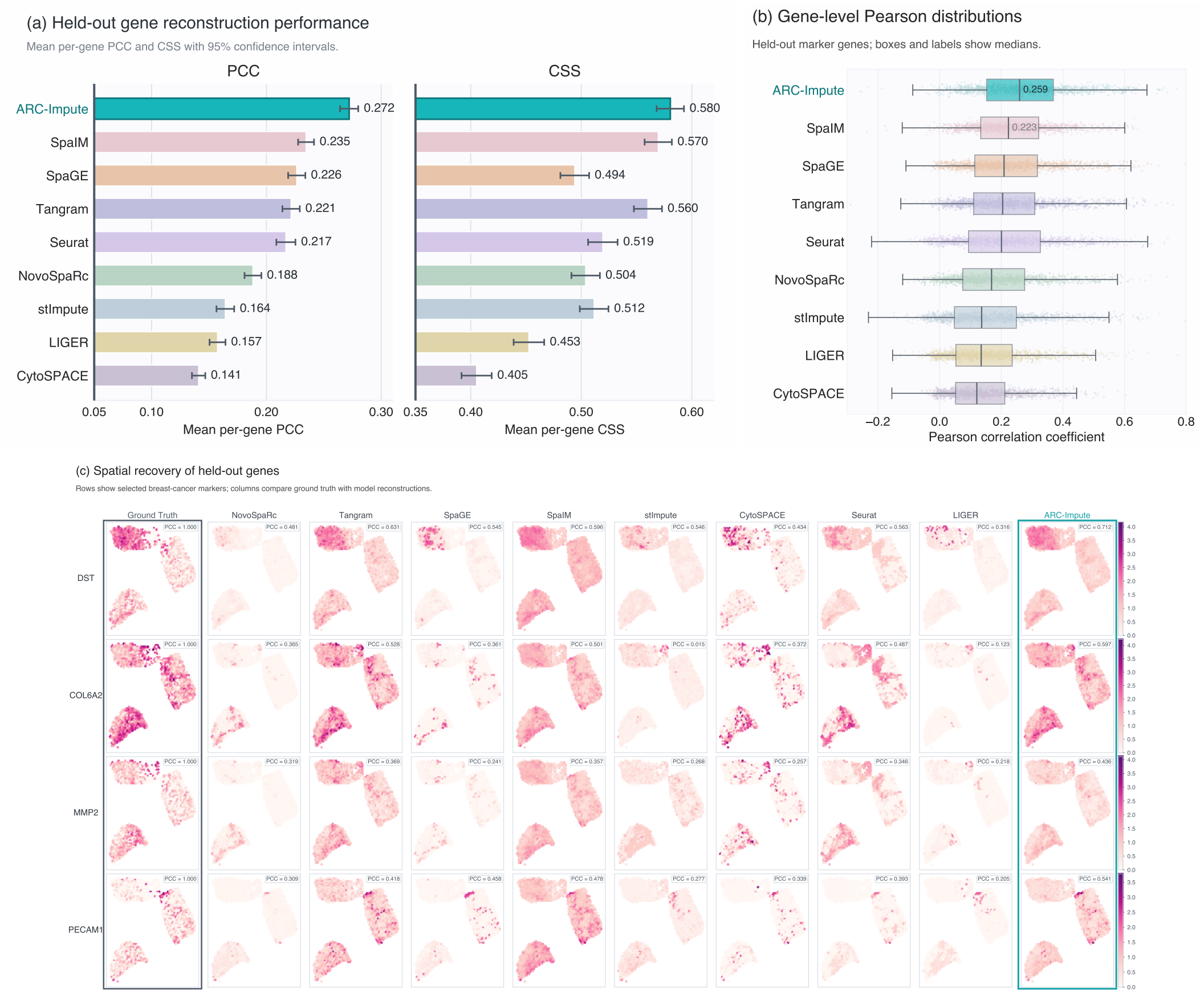
ARC-Impute improves held-out gene reconstruction in Human Breast Cancer Visium data. **(a)** Mean per-gene PCC and CSS with 95% confidence intervals across held-out genes. **(b)** Gene-level PCC distributions showing a marked upward shift in ARC-Impute’s reconstruction accuracy. **(c)** Ground truth and model reconstructions for four held-out breast-cancer-associated genes; labels report PCC across 1,160 common spots.

### 3.2. ARC-Impute achieves consistently top-ranked imputation performance across diverse datasets

Across 17 datasets spanning human and mouse tissues and spot-based and single-cell technologies, we ranked methods within each dataset by PCC, cosine similarity and SSIM. ARC-Impute ranked first for PCC in all 17 datasets and exceeded the dataset-specific mean of the eight baseline methods by 47.3% on average. It also maintained consistently high cosine-similarity rankings across both the 10 spot-based datasets (Figure 3a) and seven single-cell spatial datasets (Figure 3b), and ranked first for SSIM in 9 datasets and within the top three in 16 of 17. Although designed for mixed-cell spot-based ST, ARC-Impute retained strong performance on native single-cell datasets, demonstrating consistent performance across tissues, species, disease states and spatial resolutions.

**Figure 3.**
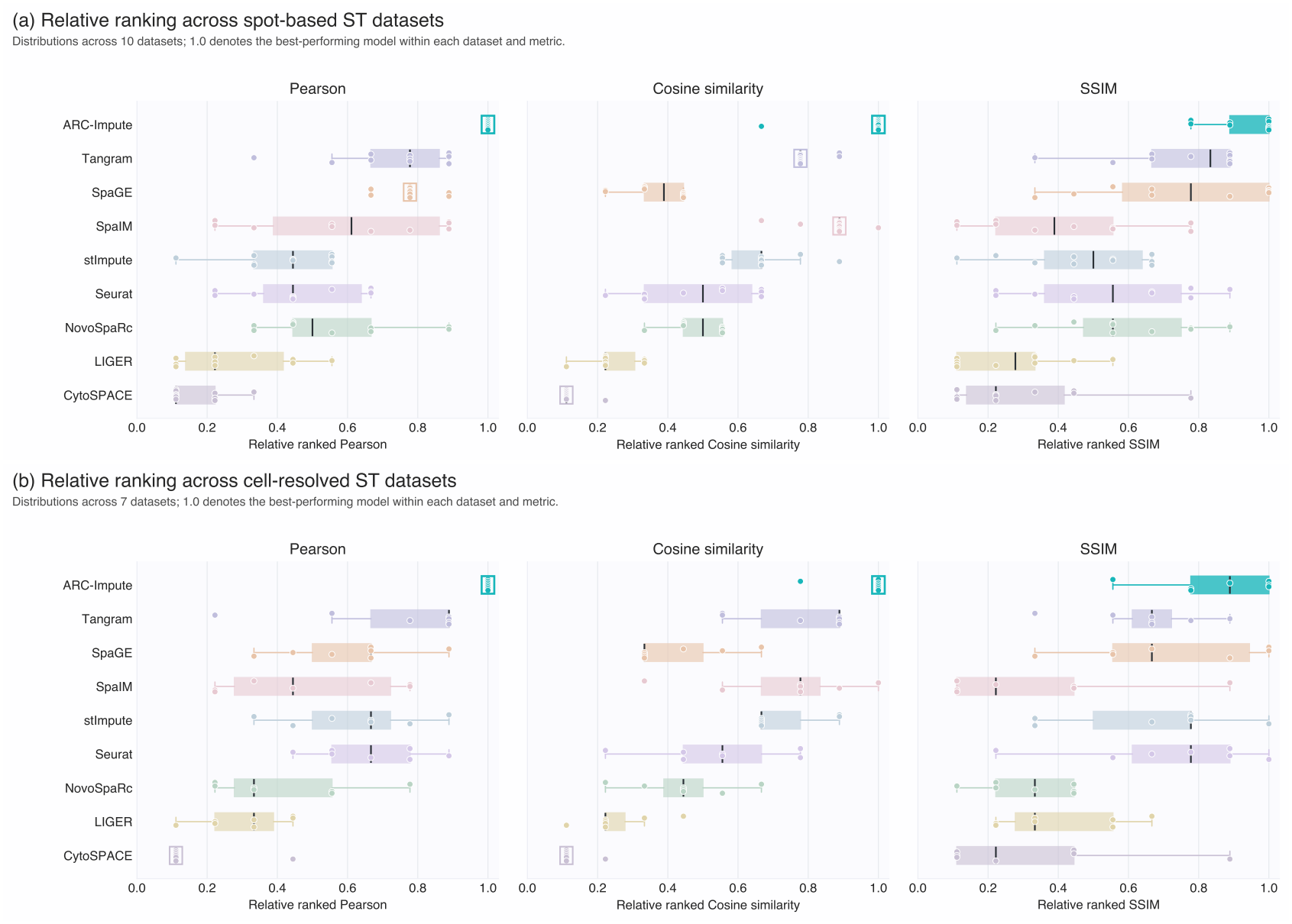
Platform-stratified reconstruction rankings across the nine-model comparison. **(a)** Ten spot-based spatial transcriptomics datasets, including three pseudospot benchmarks. **(b)** Seven native cell-resolved spatial transcriptomics datasets. Points denote individual datasets and boxes summarise their relative ranks (first place = 1.0; last place = 0.11).

### 3.3. ARC-Impute improves spot-level cell-type composition recovery

Spot-based spatial transcriptomics measures gene expression from mixtures of multiple cells, so we next assessed whether ARC-Impute could recover spot-level cell-type composition. We evaluated ARC-Impute on three pseudospot datasets with known ground-truth cell-type proportions: Allen MERFISH, osmFISH and SpaceTx ExSeq (see Supplementary Methods, Section S1.3). We compared reference-guided imputation methods that estimate spot-level cell-type composition and assessed performance using mean spot-wise Pearson correlation and cell-type RMSE. ARC-Impute performed best on both metrics across all three datasets (Figure 4a). The separation was clearest on osmFISH, where ARC-Impute achieved a spot-wise Pearson correlation of 0.573 compared with 0.344 for the next-best method, while reducing cell-type RMSE from 0.163 to 0.144.

**Figure 4.**
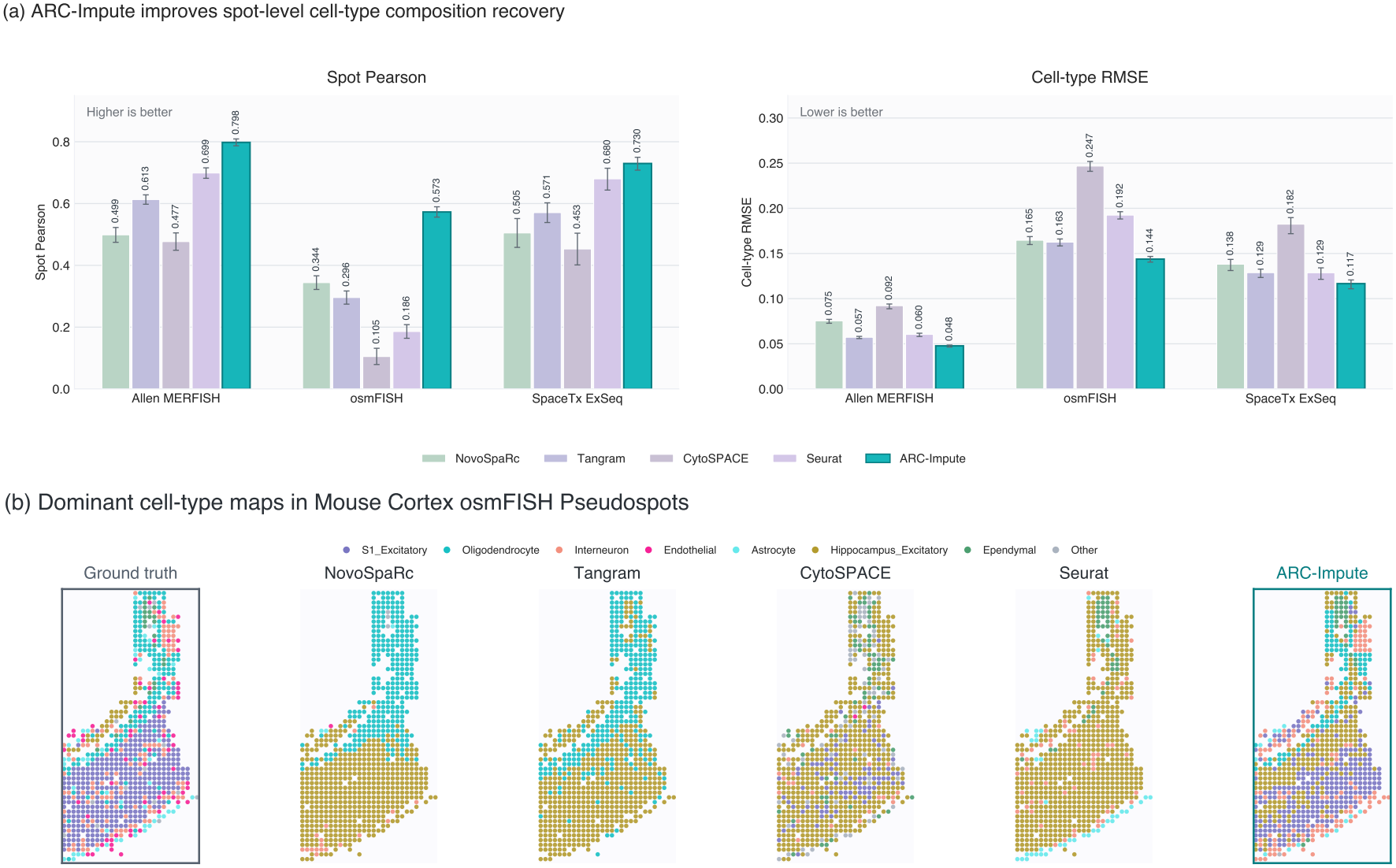
ARC-Impute improves spot-level cell-type composition recovery. **(a)** Spot-wise Pearson correlation and cell-type RMSE across three pseudospot datasets; whiskers show 95% percentile-bootstrap confidence intervals. **(b)** Ground-truth and predicted dominant cell types for osmFISH pseudospots, with rare ground-truth-dominant classes grouped as Other.

To examine how this improvement was reflected spatially, we compared predicted dominant cell-type maps in the osmFISH tissue (Figure 4b). ARC-Impute more closely recovered the broad S1 excitatory territory in the lower tissue and the oligodendrocyte-rich structure in the upper region, while retaining smaller cell populations. By contrast, several baseline methods incorrectly labelled broad territories that were S1 excitatory in the ground truth as hippocampal excitatory, or produced spatially scattered cell-type predictions. These spatial patterns were consistent with the aggregate composition metrics and suggest that ARC-Impute better preserves tissue-level cellular organisation when assigning reference cells to spatial locations.

### 3.4. ARC-Impute remains robust under reference resampling and gene-panel perturbations

Reference-guided predictions depend heavily on the reference cells and shared genes used during training. Small changes in reference composition or shared-gene selection can occur between analyses; if these lead to large changes in predictions, then the resulting biological conclusions may be unreliable (Stahl et al. 2025, Qiao and Huang 2024, Sun et al. 2024). We therefore evaluated robustness across 14 datasets using ten matched perturbation runs per dataset, with each run using 80% of the fold-training genes and a bootstrapped reference containing 80% of the cells; the spatial target and held-out genes remained fixed. ARC-Impute’s leading imputation accuracy was accompanied by the strongest prediction consistency under perturbation. For gene expression, we measured consistency as the mean pairwise PCC between the predicted spatial expression vectors for each gene across the ten runs. ARC-Impute achieved a relative-rank score of 0.902, compared with 0.848 for SpaIM, making it the only method to lead both axes of Figure 5a.

**Figure 5.**
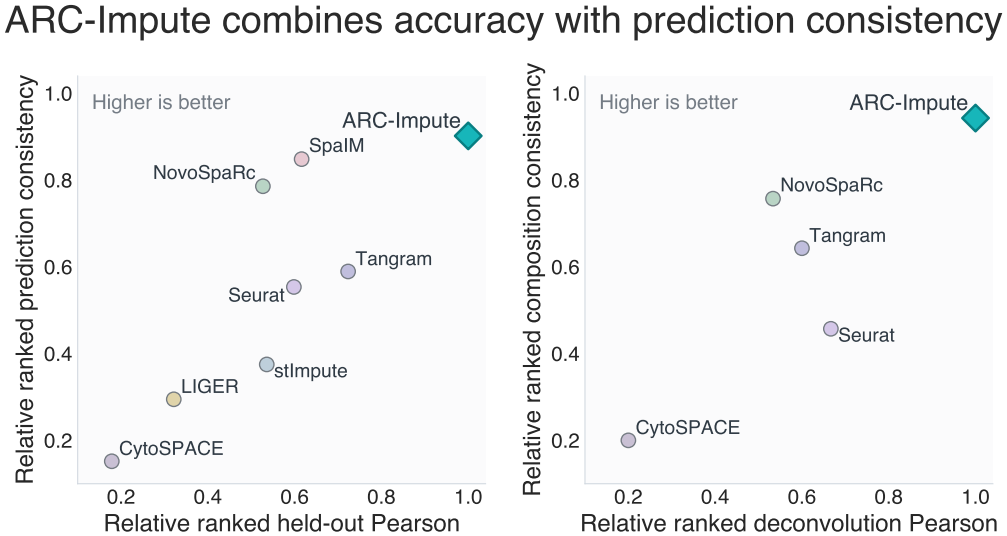
ARC-Impute combines reconstruction accuracy with prediction consistency. Mean within-dataset relative ranks, with 1.0 denoting the best-performing model. **(a)** Held-out gene reconstruction accuracy and consistency across 14 datasets. **(b)** Cell-type composition accuracy across three pseudospot benchmarks and consistency across 14 datasets.

The same pattern was observed for cell-type composition. ARC-Impute achieved the strongest cross-dataset cell-type composition consistency ranking, placing first on 11 of 14 datasets, while also ranking first for deconvolution Pearson correlation across all three ground-truth pseudospot benchmarks (Figure 5b). Its mean Jensen-Shannon divergence in cell-type composition from the ten-run consensus was 0.0285, representing reductions of 26.3% relative to NovoSpaRc and 57.4% relative to Tangram. These results indicate that ARC-Impute combines strong reconstruction and deconvolution accuracy with more reproducible predictions under variation in reference sampling and shared-gene selection.

### 3.5. Gene imputability reflects tissue-specific biological programmes

We next asked whether imputation accuracy was uniform across genes or whether some genes were easier to predict than others. For each dataset, we ranked held-out genes by their mean PCC across all nine methods (see Supplementary Methods, Section S4). This revealed a broad spread in gene-level imputation accuracy, with some genes imputed well on average across methods and others remaining difficult to predict (see Supplementary Figure S2).

Given this variation, we asked whether the more accurately imputed genes shared common biological functions. We performed gene-set enrichment analysis for each dataset, considering pathways enriched when their genes tended to rank highly by PCC-based imputation accuracy. We repeated this analysis across five human tumour datasets and five mouse neural datasets, then compared enriched pathways across datasets to determine whether similar biological programmes were repeatedly easier to impute within each biological setting.

In the five human tumour datasets, genes involved in epithelial-to-mesenchymal transition and extracellular-matrix remodelling were consistently among the most accurately imputed (Hallmark EMT: *q ≤* 0.05 in 5/5 datasets, median NES = 5.97; Reactome extracellular matrix degradation: *q ≤* 0.05 in 5/5 datasets, median NES = 4.36). These enrichments remained significant in most datasets after accounting for gene-level sparsity (4/5 datasets; median signed NES = 3.17 and 2.89, respectively), suggesting that they were not driven only by genes being broadly detected across the tissue.

The mouse neural datasets showed a different, tissue-relevant pattern. Genes involved in neuronal and synaptic function were more accurately imputed (Reactome neuronal system: *q ≤* 0.05 in 4/5 datasets, median NES = 2.32; GO synaptic signalling: *q ≤* 0.05 in 5/5 datasets, median NES = 3.56). Synaptic signalling remained enriched after accounting for gene-level sparsity (*q ≤* 0.05 in 5/5 datasets, median NES = 3.27). These results indicate that gene imputability is not random, but reflects biological programmes that are spatially structured within each tissue context.

## 4. Discussion and Conclusion

This study shows that reference-guided spatial imputation can be improved by regularising the cell-to-spot mapping itself. Because scRNA-seq and spatial transcriptomics are connected through only a limited set of shared genes, many different mappings can explain the measured genes similarly well. ARC-Impute addresses this problem by combining anchor-residual optimisation, target-guided reference curation and assignment routing, and twin-panel consistency. Together, these modules build a regularised cell-to-spot mapping that reduces local assignment ambiguity, source-target mismatch and reference-transfer instability. Across 17 benchmark datasets, ARC-Impute showed consistently strong held-out gene reconstruction across human and mouse tissues, tumour and neural datasets, and both spot-based and single-cell technologies. This breadth suggests that its benefit is not restricted to a single assay or biological setting. Rather, regularising the weakly constrained mapping appears useful across diverse forms of reference-guided spatial reconstruction.

ARC-Impute also improved spot-level cell-type composition recovery and remained stable under reference resampling and gene-panel perturbation. These results support the idea that gene imputation and cell-type composition depend on the same inferred mapping of reference cells to spatial locations. By stabilising this mapping, ARC-Impute improves both predicted expression and the reproducibility of downstream cellular interpretation.

Gene-level imputability was also structured by biological context rather than varying uniformly across genes. Extracellular-matrix and epithelial-to-mesenchymal-transition programmes were more accurately reconstructed in human tumour datasets, whereas neuronal and synaptic programmes showed greater reconstructability in mouse neural tissues. However, some highly reconstructable programmes also showed substantial variation across methods. These findings suggest that imputation reliability is biologically dependent, but still sensitive to the reconstruction method used.

ARC-Impute nevertheless inherits limitations shared by all reference-guided approaches. Its predictions remain bounded by the coverage and quality of the scRNA-seq reference, and cell states or expression programmes absent from the reference cannot be directly recovered. Evaluation also remains challenging because held-out measured genes provide only a proxy for truly unmeasured expression, while constructed pseudospots do not fully reproduce native spot capture. Future work should explore multi-reference integration, detection of unsupported target states and validation using paired or orthogonal spatial measurements. In summary, ARC-Impute provides a more accurate and reproducible approach to reference-guided spatial reconstruction across diverse biological and technical settings.

## 5. Author contributions

JY, JK, EH jointly conceived and led the conceptual framework of the study with design and implementation input from AZ. EH led the method development and implementation and evaluation of the framework, with strong input from JK, JY and AZ. EH, JY and JK led the initial draft with input from AZ with all authors contributing to the writing, editing, and approval of the manuscript.

## 6. Supplementary material

Supplementary material accompanies this preprint.

## 9. Data availability

Dataset accessions and persistent identifiers are provided in Supplementary Figure S1. ARC-Impute source code, documentation, and plotting data underlying Figures 2–5 are archived on Zenodo (v0.1.0; https://doi.org/10.5281/zenodo.22722992), with the GitHub repository available at https://github.com/ethanhaan/arc-impute.

