## Supplementary material for "ARC-Impute: robust reference-guided spatial gene imputation through regularised cell-to-spot mapping"

### S1. Datasets, references, preprocessing and pseudospot construction

#### S1.1. Dataset selection and reference pairing

The benchmark comprised 17 spatial target–reference configurations: seven native spot-level targets, three pseudospot targets derived from cell-resolved assays and seven native cell-resolved targets (Supplementary Figure S1). The human tumour sources and references are described by Moncada et al. (2020), Andersson et al. (2021), Wu et al. (2021), Song et al. (2022), Arora et al. (2023), Kurkalang et al. (2023); the mouse neural sources and references by Zeisel et al. (2015), Ståhl et al. (2016), Codeluppi et al. (2018), Tasic et al. (2018), Tepe et al. (2018), Wang et al. (2018), Eng et al. (2019), Yao et al. (2023), Zhang et al. (2023).

Same-patient references were used for CID44971 breast cancer and PDAC-A. Where a same-patient reference was unavailable, a pairing already used in a published analysis was retained: HER2ST was paired with the five-donor HER2-positive Wu et al. reference (Andersson et al. 2021); mouse olfactory-bulb replicate 12 with GSE121891 (Liu et al. 2024); osmFISH with the Zeisel mouse-brain reference (Abdelaal et al. 2020); and the SpaceTx VISp assays with the Tasic 2018 Allen VISp reference (Zhang et al. 2023). The remaining external references were selected by species, anatomical region and disease context. Supplementary Figure S1 gives the complete source identifiers and pairings, processed dimensions and annotation fields.

| Spatial target | Spatial source / accession | Spatial context and assay | scRNA-seq reference | Reference source / accession | Benchmark dimensions | Labels used |
|---|---|---|---|---|---|---|
| Spot-based targets (n = 10) | | | | | | |
| Breast cancer Visium CID44971 | Zenodo 4739739; GSE176078 | Human; breast; TNBC 10x Visium; spot | CID44971 same-patient scRNA-seq | GSE176078; GSM5354531 | 1,160 spots \| 7,986 ref. cells 1,475 shared genes | Markers: celltype_minor ARC: Classification (author pathology) |
| Mouse cortex Visium Cortex_1-4 | 10x V1 Adult Mouse Brain; Squidpy dataset | Mouse; cortex; healthy 10x Visium; spot | Tasic/Allen mouse cortex scRNA-seq | Squidpy sc_mouse_cortex; Tasic 2018 | 949 spots \| 21,697 ref. cells 1,293 shared genes | Markers: cell_subclass ARC: cluster (Squidpy/Allen) |
| PDAC-A ST1 GSM3036911 | GSE111672; GSM3036911 | Human; pancreas; PDAC legacy ST; spot | PDAC-A same-patient inDrop scRNA-seq | GSE111672 | 426 spots \| 1,926 ref. cells 1,286 shared genes | Markers: celltype ARC: layer_manual_PDAC (pathology) |
| HER2ST B1 | HER2ST; DOI 10.1038/s41467-021-26271-2 | Human; breast; HER2+ cancer legacy ST; spot | Five HER2+ breast cancer donors | GSE176078 | 268 spots \| 4,226 ref. cells 655 shared genes | Markers: celltype_major ARC: author pathologist labels |
| Mouse olfactory bulb Rep12 | Stahl ST Rep12; SDePER annotation | Mouse; olfactory bulb; healthy legacy ST; spot | WT1 + WT2 mouse olfactory bulb scRNA-seq | GSE121891 | 278 spots \| 4,558 ref. cells 421 shared genes | Markers: celltype ARC: SDePER anatomical annotation |
| OSCC sample 12 GSM6339642 | GSE208253; GSM6339642 | Human; oral; HPV- OSCC 10x Visium; spot | OSCC scRNA-seq cohort | GSE215403 | 1,437 spots \| 28,186 ref. cells 634 shared genes | Markers: cell_type ARC: pathologist_anno.x |
| Prostate cancer FFPE | 10x Visium CytAssist FFPE prostate | Human; prostate; invasive cancer 10x Visium FFPE; spot | Human prostate scRNA-seq cohort | GSE176031 | 4,321 spots \| 21,743 ref. cells 861 shared genes | Markers: ID ARC: CellPie/10x histology label |
| Allen MERFISH Zhuang-ABCA-2.037 Pseudospot | Allen ABC Atlas; Zhuang-ABCA-2.037 | Mouse; brain; healthy MERFISH; pseudospot | Allen whole-mouse-brain 10x reference | Allen ABC Atlas; WMB-10Xv3 | 674 pseudospots \| 22,095 ref. cells 300 shared genes | Markers: subclass-derived cell_type ARC: Allen region_layer; majority vote |
| osmFISH cortex Pseudospot | osmFISH; DOI 10.1038/s41592-018-0175-z | Mouse; cortex; healthy osmFISH; pseudospot | Zeisel mouse-brain scRNA-seq reference | ZeiselBrainData; DOI 10.1126/science.aaa1934 | 794 pseudospots \| 2,816 ref. cells 33 shared genes | Markers: level2class-derived cell_type ARC: author Region; majority vote |
| SpaceTx ExSeq VISp Pseudospot | SpaceTx ExSeq; DOI 10.1038/s41598-023-36638-8 | Mouse; VISp; healthy ExSeq; pseudospot | Allen VISp Smart-seq shared subclasses | Allen VISp Smart-seq; archive 694413985 | 203 pseudospots \| 13,523 ref. cells 42 shared genes | Markers: cell_type ARC: SpaceTx layer; majority vote |
| Native cell-resolved targets (n = 7) | | | | | | |
| Allen MERFISH Zhuang-ABCA-2.037 | Allen ABC Atlas; Zhuang-ABCA-2.037 | Mouse; brain; healthy MERFISH; native cell | Allen whole-mouse-brain 10x reference | Allen ABC Atlas; WMB-10Xv3 | 4,081 cells \| 22,095 ref. cells 300 shared genes | Markers: subclass-derived cell_type ARC: Allen region_layer |
| osmFISH somatosensory cortex | osmFISH; DOI 10.1038/s41592-018-0175-z | Mouse; cortex; healthy osmFISH; native cell | Zeisel mouse-brain scRNA-seq reference | ZeiselBrainData; DOI 10.1126/science.aaa1934 | 4,839 cells \| 2,816 ref. cells 33 shared genes | Markers: level2class-derived cell_type ARC: author Region |
| STARmap visual cortex 20180505_BY3_1kgenes | STARmap; Zenodo 10698912 | Mouse; visual cortex; healthy STARmap; native cell | Allen whole-mouse-brain 10x reference | Allen ABC Atlas; WMB-10Xv3 | 1,207 cells \| 22,095 ref. cells 375 shared genes | Markers: cell_type ARC: seven author anatomical labels |
| seqFISH+ cortex/SVZ FOV0-6 | seqFISH+; DOI 10.1038/s41586-019-1049-y | Mouse; cortex/SVZ; healthy seqFISH+; native cell | Allen whole-mouse-brain 10x reference | Allen ABC Atlas; WMB-10Xv3 | 913 cells \| 22,095 ref. cells 1,046 shared genes | Markers: cell_type ARC: author Region |
| SpaceTx ISS VISp | SpaceTx ISS; DOI 10.1038/s41598-023-36638-8 | Mouse; VISp; healthy ISS; native cell | Allen VISp Smart-seq reference | Allen VISp Smart-seq; archive 694413985 | 3,065 cells \| 14,249 ref. cells 119 shared genes | Markers: cell_type ARC: SpaceTx layer |
| SpaceTx BARISTASeq VISp | SpaceTx BARISTASeq; DOI 10.1038/s41598-023-36638-8 | Mouse; VISp; healthy BARISTASeq; native cell | Allen VISp Smart-seq reference | Allen VISp Smart-seq; archive 694413985 | 4,432 cells \| 14,249 ref. cells 76 shared genes | Markers: cell_type ARC: SpaceTx layer |
| SpaceTx ExSeq VISp | SpaceTx ExSeq; DOI 10.1038/s41598-023-36638-8 | Mouse; VISp; healthy ExSeq; native cell | Allen VISp Smart-seq reference | Allen VISp Smart-seq; archive 694413985 | 1,240 cells \| 14,249 ref. cells 42 shared genes | Markers: cell_type ARC: SpaceTx layer |

**Figure S1** Dataset and reference inventory for the 17 benchmark configurations. Rows report each spatial target, source and assay, paired scRNA-seq reference, processed benchmark dimensions, and annotation fields used for marker selection and ARC-Impute regional anchors.

#### S1.2. Preprocessing and annotations

Source count matrices and published annotations were retained wherever possible. Raw counts were stored alongside a transformed analysis representation. This representation was independently library-size normalised to 10,000 counts per observation and log-transformed ($log(1+x)$). The exception was the library-provided Squidpy mouse-cortex pair, which was already supplied in median-library-size-normalised and log-transformed form. Each model received either raw counts or transformed expression according to its input requirements, as detailed in Supplementary Table S1. Source-specific quality control was also retained: osmFISH cells were restricted to the authors' valid-cell flag, and the prepared SpaceTx ExSeq targets excluded 31 cells annotated as outside VISp. Gene names were made unique before the reference and target were restricted to their shared evaluation panel.

Marker-gene groups retained the biological resolution supplied by each reference, including cell type, subclass, major cell type or source identifier, rather than imposing a new cross-dataset ontology. ARC-Impute region anchors preferentially used author-provided pathology or anatomical labels. Derived labels were used only where preparation explicitly collapsed a published hierarchy, such as Allen cortical layers, or where a published third-party anatomical annotation was supplied. The exact marker and region fields are reported in Supplementary Figure S1.

#### S1.3. Pseudospot construction

Allen MERFISH, osmFISH and SpaceTx ExSeq cells were aggregated into deterministic square spatial bins. For each dataset, the grid width was selected so that retained bins contained approximately six cells on average; bins containing fewer than two cells were discarded. Raw expression counts were summed across constituent cells before normalisation, and the geometric centre of each bin was used as its pseudospot coordinate. The pseudospot cell-type composition for class $c$ was the number of constituent cells assigned to $c$ divided by the total number of retained cells in that bin. Dominant cell-type and region labels were assigned by deterministic majority vote.

For SpaceTx ExSeq, cell types were the released geometric-median consensus strategy assignments harmonised to broad Tasic 2018 subclasses. These are consensus-derived benchmark labels rather than independently measured biological ground truth. For the osmFISH maps in Figure 4(b), a cell type was displayed separately only if it was dominant in more than five ground-truth pseudospots. Classes dominant in five or fewer ground-truth pseudospots were grouped as *Other*, and the same grouping was then applied to every prediction panel so that colours remained directly comparable.

### S2. Benchmark design, evaluation, uncertainty and robustness

Where results from the reconstruction, cell-type-composition and reference-transfer evaluations were summarised across datasets, scores were first converted to within-dataset relative ranks. For method $m$, metric $q$ and dataset $d$, the relative rank was

$$R_{d,m,q} = \frac{N_{d,q} - r_{d,m,q} + 1}{N_{d,q}},$$

where $N_{d,q}$ is the number of eligible methods and $r_{d,m,q}$ is the conventional rank, with rank 1 assigned to the best result. Higher values were better for correlation, cosine similarity, SSIM and expression-prediction consistency; lower values were better for RMSE and Jensen–Shannon divergence. Ties received their average rank. First place therefore received 1.0, while ninth place in the nine-method reconstruction comparison received $1/9 = 0.11$. Cross-dataset summaries are unweighted means of the relative ranks.

#### S2.1. Held-out reconstruction benchmark

**Gene selection and cross-validation folds.** Evaluation genes were restricted to those shared by each spatial target and its paired scRNA-seq reference. For most datasets, the panel was the deduplicated union of the top 100 genes ranked within each annotated reference group, intersected with the shared genes. When overlap was too limited, all shared genes were retained. The panel was divided into five shuffled folds using `random_state=1`. Each fold held out approximately 20% of the genes and used the remaining genes for training. Every selected gene was held out once, and all methods were evaluated on the same genes. Held-out genes were never supplied during model preparation or fitting.

Let $y_{s,g}$ and $\hat{y}_{s,g}$ denote the observed and predicted expression of held-out gene $g$ at spatial observation $s$, respectively, and let $S$ be the number of evaluated spatial observations.

**CSS.** Cosine similarity measured agreement in the direction of the observed and predicted expression vectors for each gene:

$$\mathrm{CSS}_g = \frac{\sum_{s=1}^{S} y_{s,g}\hat{y}_{s,g}}{\sqrt{\sum_{s=1}^{S} y_{s,g}^2}\sqrt{\sum_{s=1}^{S} \hat{y}_{s,g}^2}}.$$

**SSIM.** Structural similarity compared the level, variance and covariance of the observed and predicted spatial expression vectors. Each vector was independently divided by its maximum to give $\tilde{y}_g$ and $\tilde{\hat{y}}_g$:

$$\mathrm{SSIM}_g = \frac{(2\mu_y\mu_{\hat{y}} + C_1)(2\sigma_{y\hat{y}} + C_2)}{(\mu_y^2 + \mu_{\hat{y}}^2 + C_1)(\sigma_y^2 + \sigma_{\hat{y}}^2 + C_2)},$$

where $\mu_y$ and $\mu_{\hat{y}}$ are the means, $\sigma_y^2$ and $\sigma_{\hat{y}}^2$ are the population variances, and $\sigma_{y\hat{y}}$ is the covariance of the scaled vectors. The constants were $C_1 = (0.01M)^2$ and $C_2 = (0.03M)^2$, where $M$ is the maximum dynamic range, normally 1. SSIM was applied once to each gene's expression vectors across all spatial observations. It did not use coordinates, rasterisation, sliding windows or spatial-neighbour weights. Undefined entries caused by a zero maximum were replaced by $10^{-20}$, matching the reported implementation.

**PCC.** Pearson correlation measured linear agreement in spatial expression for each gene:

$$r_g = \frac{\sum_{s=1}^{S}(y_{s,g} - \bar{y}_g)(\hat{y}_{s,g} - \bar{\hat{y}}_g)}{\sqrt{\sum_{s=1}^{S}(y_{s,g} - \bar{y}_g)^2}\sqrt{\sum_{s=1}^{S}(\hat{y}_{s,g} - \bar{\hat{y}}_g)^2}}.$$

**Aggregation and exclusions.** Per-gene values from the five test folds were pooled within each dataset and averaged, so every held-out gene contributed once. PCC was non-evaluable when either vector was constant, and CSS was non-evaluable when either vector had zero norm. These results were stored as missing and omitted from the corresponding mean. Only completed, shape-consistent outputs containing finite predictions entered the summaries; failed runs remained explicit error records in Supplementary Table S1.

**Figure 2 uncertainty.** The 95% intervals in Figure 2(a) were calculated across non-missing held-out-gene values as $1.96s_g/\sqrt{n_g}$, where $s_g$ is the sample standard deviation and $n_g$ is the number of evaluable genes. They describe variation across held-out genes, not repeated model fits. In Figure 2(b), boxes show the median and interquartile range; whiskers extend to the most extreme values within 1.5 times the interquartile range.

#### S2.2. Cell-type composition evaluation

**Evaluation cohort.** Composition accuracy was evaluated on the Allen MERFISH, osmFISH and SpaceTx ExSeq pseudospot targets, which had known spot-level cell-type proportions. Ground-truth and predicted compositions were represented as cell-type proportion vectors for each spot, normalised to sum to one. NovoSpaRc, Tangram, CytoSPACE, Seurat and ARC-Impute were included because they produced spot-level compositions. SpaIM, stImpute, LIGER and SpaGE were excluded because they do not natively support cell-type deconvolution and did not produce calibrated spot-level mixtures.

**Spot-wise PCC.** Let $t_{s,c}$ and $p_{s,c}$ denote the true and predicted proportions of cell type $c$ in spot $s$, and let $S$ and $C$ be the numbers of spots and cell types. PCC was calculated between the true and predicted composition vectors for each spot:

$$r_s = \frac{\sum_{c=1}^{C}(t_{s,c} - \bar{t}_s)(p_{s,c} - \bar{p}_s)}{\sqrt{\sum_{c=1}^{C}(t_{s,c} - \bar{t}_s)^2}\sqrt{\sum_{c=1}^{C}(p_{s,c} - \bar{p}_s)^2}}.$$

The reported value was the mean across evaluable spots. Spots with a constant true or predicted vector were omitted.

**Cell-type RMSE.** RMSE was calculated across spots for each cell type:

$$\mathrm{RMSE}_c = \sqrt{\frac{1}{S}\sum_{s=1}^{S}(p_{s,c} - t_{s,c})^2}, \qquad \overline{\mathrm{RMSE}} = \frac{1}{C}\sum_{c=1}^{C}\mathrm{RMSE}_c.$$

The reported value was the unweighted mean across cell types.

**Figure 4 uncertainty.** Fitted predictions remained fixed while paired true and predicted spot rows were resampled with replacement. Spot-wise PCC and cell-type RMSE were recomputed for 10,000 bootstrap samples, and the 2.5th and 97.5th percentiles formed the 95% interval. These resamples quantify spot-level uncertainty, not repeated model training.

#### S2.3. Reference-transfer stability

**Matched perturbation design.** Stability was evaluated for 14 native benchmark targets: seven measured spot-level datasets and seven native cell-resolved datasets. The three pseudospot derivatives were excluded. Each dataset had ten matched runs using seeds 1–10. The spatial target and held-out test genes remained fixed. In each run, 80% of the fold-training genes were sampled without replacement, and the scRNA-seq reference was replaced by a bootstrap sample containing 80% of the original number of cells. The same seed controlled both perturbations. Expression-prediction consistency was evaluated for NovoSpaRc, Tangram, SpaIM, stImpute, CytoSPACE, Seurat, LIGER and ARC-Impute. Composition consistency was evaluated for the five composition-eligible methods.

**Expression-prediction consistency.** Analysis was restricted to spatial observations present in all ten runs. For each held-out gene $g$, PCC was calculated for every pair of run-specific prediction vectors and averaged:

$$\mathrm{PC}_g = \frac{1}{|\mathcal{P}_g|} \sum_{(a,b)\in\mathcal{P}_g} \mathrm{PCC}(\hat{\mathbf{y}}_g^{(a)}, \hat{\mathbf{y}}_g^{(b)}), \qquad K = 10,$$

where $\mathcal{P}_g$ is the set of evaluable pairs among the 45 possible run pairs. Pairs containing a constant prediction vector were omitted. The dataset score was the mean across evaluable genes.

**Cell-type-composition consistency.** For composition-capable methods, each spot in each run was represented by a vector of predicted cell-type proportions, normalised to sum to one. The mean of the ten run-specific vectors, also normalised to sum to one, defined the consensus composition $\bar{\mathbf{p}}_s$ for that spot. The divergence of run $k$ from this consensus at spot $s$ was

$$\mathrm{JSD}(\mathbf{p}_s^{(k)}, \bar{\mathbf{p}}_s) = \frac{1}{2}\mathrm{KL}(\mathbf{p}_s^{(k)} \,\|\, \mathbf{m}_s^{(k)}) + \frac{1}{2}\mathrm{KL}(\bar{\mathbf{p}}_s \,\|\, \mathbf{m}_s^{(k)}), \qquad \mathbf{m}_s^{(k)} = \frac{\mathbf{p}_s^{(k)} + \bar{\mathbf{p}}_s}{2}.$$

JSD was averaged across spots for each run and then across the ten runs. Lower mean JSD indicates greater cell-type-composition consistency and therefore received the better within-dataset rank.

### S3. Model provenance, implementation and reproducibility

**Table S1.** Model implementation and reproducibility. Software provenance, method-specific inputs and preprocessing, benchmark adaptations and fit settings for the nine compared methods.

| Method | Software provenance | Inputs, preprocessing and adaptations | Fit settings |
|---|---|---|---|
| **NovoSpaRc** | `github.com/rajewsky-lab/novosparc`; v0.4.4 official wheel. | Log-normalised fold data were supplied. NovoSpaRc's default 5-nearest-neighbour source and spatial graphs were retained. The initial entropy regularisation was increased from $5 \times 10^{-4}$ to $5 \times 10^{-3}$ for numerical stability. | $\alpha = 0.5$; initial $\epsilon = 5 \times 10^{-3}$; native tenfold $\epsilon$ search for numerical failures; seed 0; target-marginal scaling. |
| **Tangram** | `github.com/broadinstitute/Tangram`; tangram-sc v1.0.4, tag commit `d537ff9`. | Tangram's standard log-normalised input and cell-level mapping were retained. Fits received fold-training genes only and used the RNA-count-based spot-density prior. | Adam; learning rate 0.1; 500 epochs; seed 0; official reconstruction and density objective. |
| **SpaIM** | `github.com/QSong-github/SpaIM`; upstream snapshot `3d8b76b`. | Following SpaIM's clustered-reference design, reference cells were aggregated into Leiden mean profiles (resolution 0.5), with the official fourfold augmentation and 50% random zero masking retained. Benchmark inputs were library-normalised before log transformation rather than passed through the upstream log-only loader; upstream expression filtering was not repeated. | Adam, learning rate $10^{-3}$; hidden widths 256/512; batch 500; 200 epochs; seed 1234. |
| **stImpute** | `github.com/cquzys/stImpute`; upstream commit `5a09659`. | Official log-normalised inputs and ESM-2 gene embeddings were used. Genes without embeddings used stImpute's expression-similarity graph rather than being excluded. | Adam, learning rate $10^{-2}$; 30 autoencoder epochs, two EM cycles and the official size-adaptive 50/20-neighbour schedule; seed 42. |
| **CytoSPACE** | `github.com/digitalcytometry/cytospace`; v1.1.0, upstream commit `bb7fe12`. | CytoSPACE's official raw-count assignment, internal CPM plus log2 transformation and automatic Seurat cell-fraction estimation were retained. Smart-seq references used the default 1,500-transcript downsampling. Cells and spots with zero fold-training counts were removed. | Pearson cost; `lapjv` assignment; mean 5 cells per spot, including legacy ST; one process; seed 1. |
| **Seurat** | `github.com/satijalab/seurat`; Seurat v5.3.1, tag commit `ca0ab0f`. | Seurat's official raw-count SCT and anchor-transfer workflow was used. PCA dimensions and anchor weights were capped only where required by fold size. Zero-support reference cells were removed; zero-support spatial observations were restored using reference-mean predictions and reference-frequency compositions. | Up to 30 PCs; all available dimensions; $k.\mathrm{weight} \le 50$; seed 1; official anchor transfer. |
| **LIGER** | `github.com/welch-lab/pyliger`; PyLiger v0.2.4, tag commit `45b40e6`. | LIGER's standard raw-count library normalisation, scaling without centring, iNMF and quantile normalisation were used with fold-specific shared genes. Four zero-support spatial observations across Allen MERFISH, osmFISH and SpaceTx ISS were excluded. | $k \le 30$; $\lambda = 5$; one ALS replicate; 30 iterations; 50-nearest-neighbour transfer; seed 1. |
| **SpaGE** | `github.com/tabdelaal/SpaGE`; upstream commit `edd0fb7`. | SpaGE's standard log-normalised input and principal-vector alignment were used. Alignment was restricted to fold-training genes to prevent held-out-gene leakage. | Up to 30 principal vectors; 50-nearest-neighbour prediction; seed 0; official SpaGE alignment and transfer. |
| **ARC-Impute** | `github.com/ethanhaan/arc-impute`. | The public ARC-Impute workflow used log-normalised fold data, dataset-provided region labels, centroid source-blend reference curation (50% budget, minimum 32 per label), a persistent regional anchor and twin-loss panels. | Adam, learning rate 0.1; 500 epochs; seed 0; anchor strength 0.5; anchor KL 0.03; gate L1 0.01; twin agreement 0.05. |

**Compute environment.** All methods were run on a single NVIDIA GeForce RTX 3090 GPU (24 GB VRAM) with an Intel Core i9-9900K CPU and 64 GB system RAM.

**Shared benchmark policy.** Each cross-validation fit saw only its training-gene panel; held-out genes were used only for prediction and evaluation. Repeated perturbation runs used seeds 1–10. Only completed, shape-consistent finite outputs entered summaries; failed folds were retained as explicit error records and were not silently replaced. **Composition eligibility.** NovoSpaRc, Tangram, CytoSPACE, Seurat and ARC-Impute returned spot-level cell-type compositions. SpaIM, stImpute, LIGER and SpaGE were excluded from composition evaluation because their benchmark outputs are not calibrated spot-level mixtures.

### S4. Gene-program enrichment analysis and supporting results

This section provides a concise, method-general account of how biological programmes were tested across five human tumour and five mouse neural datasets using the same nine-method comparison as the reconstruction benchmark.

##### S4.1. Gene rankings and enrichment testing

Within each dataset, gene-set enrichment analysis (GSEA) tested whether members of a predefined biological programme accumulated near either end of a gene ranking (Subramanian et al. 2005). The signed enrichment score (ES) was the largest absolute deviation of the running score, and the normalised enrichment score (NES) placed it relative to gene sets of comparable size. Here, enrichment denotes reconstruction performance, not biological pathway activation or suppression.

Genes were ranked by mean held-out PCC across the nine methods, sparsity-adjusted mean PCC or population standard deviation across methods. Positive NES indicated higher reconstructability for the first two rankings or greater between-method instability for the third; negative NES indicated the corresponding lower-ranking end. The resulting gene-level PCC distributions and representative per-method values are shown in Supplementary Figure S2.

Signed, bidirectional GSEA used `fgseaMultilevel` with rank-only weights (`gseaParam=0`), so only gene order influenced the running score (Korotkevich et al. 2019). Parameters were seed 1729, `sampleSize=101`, `nPermSimple=10,000` and $\epsilon = 10^{-50}$. Benjamini–Hochberg correction was applied within each prespecified analysis tier, and tier-adjusted $q \leq 0.05$ defined significance. Invalid PCC cases were handled as follows: constant spatial truth was non-evaluable; a constant finite prediction was assigned PCC zero when truth varied and its error was finite; all other unexplained non-finite values failed validation.

##### S4.2. Gene-set mapping, analysis background and programme coverage

Gene programmes came from MSigDB 2025.1: Hallmark and Reactome formed the primary tier and GO Biological Process the supplementary tier (Liberzon et al. 2011). Human and mouse datasets used species-native collections and were analysed separately. Symbols were matched case-insensitively; unresolved identifiers were converted through g:Profiler and accepted only if the result occurred in the relevant GMT file. Mappings and GMT files were then fixed before testing.

The ranked background contained every evaluated gene with a finite value; unrecognised genes were retained under miss-only identifiers so that the background was not restricted to programme members. Across datasets, 67.3–93.3% of genes were recognised and effective backgrounds contained 979–3,000 genes. Programmes required 15–500 covered members, an established GSEA window limiting unstable normalisation from very small sets and overly broad results (Subramanian et al. 2005); the minimum was 10 only for the smaller targeted Allen MERFISH and STARmap panels.

Displayed programmes covered 45–307 ranked genes, of which 11–192 formed the leading edge contributing most strongly to the enrichment score. Illustrative recurrent genes included *COL1A2* and *FN1* for EMT and extracellular matrix, *PLAU* for KRAS and TNF$\alpha$/NF-$\kappa$B, and *Slc17a7* for neuronal and synaptic programmes. Inference remained programme-level.

##### S4.3. Sparsity adjustment and sensitivity analysis

Spatial sparsity can constrain gene prediction in spatial transcriptomics (Biancalani et al. 2021, Zhang et al. 2022, Wang et al. 2025). We defined it as each gene's observed spatial zero fraction and fitted mean PCC using a dataset-specific cubic regression spline with five quantile-spaced knots. The adjusted score was the residual: observed mean PCC minus its fitted expectation. Five-fold cross-validation over genes, shuffled with seed 1729, showed moderate but incomplete explanatory strength (median $R^2 = 0.43$, range 0.26–0.74). Across datasets, the median sparsity–PCC Spearman correlation was $-0.65$ (range $-0.84$ to $-0.47$), while raw and adjusted gene ranks retained a median correlation of 0.66 (range 0.33–0.83). We therefore treated the adjusted ranking as a sensitivity analysis rather than a replacement for mean PCC; the procedure did not assume that every zero was a technical dropout.

The main conclusions were largely retained after adjustment. EMT and extracellular-matrix programmes remained significantly enriched toward higher reconstruction performance in four of five human datasets, while KRAS signalling, TNF$\alpha$/NF-$\kappa$B signalling and mouse synaptic signalling remained enriched in all five corresponding datasets. Neuronal-system enrichment was less robust after adjustment, reaching significance in two of five mouse datasets.

##### S4.4. Cross-dataset recurrence and supporting results

Recurrence was the number of same-species datasets with tier-adjusted $q \leq 0.05$ in the same direction, out of the five datasets in which each programme was tested; human and mouse results were never pooled. Table S2 reports the median signed NES across all five tested datasets, so a contrary direction remains represented rather than being discarded. Before adjustment, EMT and ECM degradation recurred in all five human datasets, while neuronal-system and synaptic-signalling enrichment recurred in four and five mouse datasets, respectively.

Between-method variability was interpreted separately from average reconstruction performance. EMT and ECM organisation were enriched toward higher variability in all five human datasets, and synaptic signalling in four of five mouse datasets. This pattern was not universal: neuronal-system variability enrichment reached significance in only two mouse datasets, and none of the highlighted programmes showed significant enrichment toward lower variability. Thus, higher mean performance did not by itself imply either agreement or disagreement among methods. Complete programme-level results are reported in Table S2.

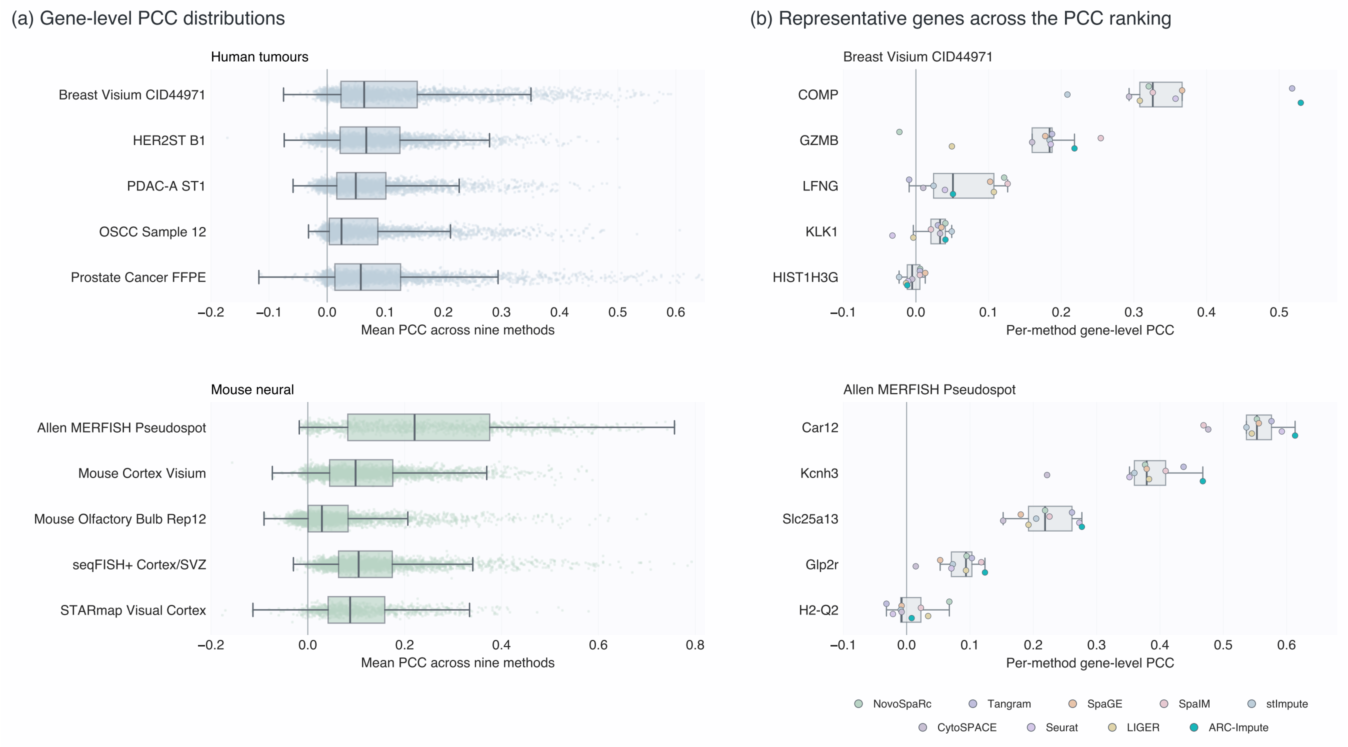


**Figure S2 Gene-level imputation accuracy varies across genes and methods. (a)** Distributions of each held-out gene's mean PCC across the nine reconstruction methods for five human tumour and five mouse neural datasets. Points represent genes; boxes show the median and interquartile range, with whiskers extending to the most extreme value within 1.5 times the interquartile range. **(b)** Method-specific PCC values for genes nearest the 5th, 25th, 50th, 75th and 95th percentiles of mean PCC in Breast Visium CID44971 and Allen MERFISH Pseudospot. Each coloured point represents one method and each box summarises the nine values. Separate horizontal scales are used for the two illustrative datasets.

**Table S2.** Gene-program enrichment across the fixed ten-dataset cohort. Cells report signed normalised enrichment scores (NES): teal denotes positive enrichment, coral denotes negative enrichment and colour intensity increases with magnitude. Bold values marked with an asterisk have tier-adjusted $q \leq 0.05$. The $k/5$ column gives the number of significant same-species datasets in the displayed high direction; median NES is the median signed value across all five tested datasets.

**Human tumour datasets**

| Programme | Breast Visium CID44971 | HER2ST B1 | PDAC-A ST1 | OSCC Sample 12 | Prostate Cancer FFPE | $k/5$ | Median NES |
|---|---|---|---|---|---|---|---|
| **Mean PCC** | | | | | | | |
| EMT | **6.86**$^*$ | **2.97**$^*$ | **5.83**$^*$ | **6.48**$^*$ | **5.97**$^*$ | 5/5 | 5.97 |
| ECM degradation | **4.36**$^*$ | **2.04**$^*$ | **5.84**$^*$ | **5.12**$^*$ | **3.44**$^*$ | 5/5 | 4.36 |
| **Sparsity-adjusted mean PCC** | | | | | | | |
| EMT | **4.17**$^*$ | -0.49 | **2.68**$^*$ | **4.68**$^*$ | **3.17**$^*$ | 4/5 | 3.17 |
| ECM degradation | **3.32**$^*$ | -0.47 | **2.89**$^*$ | **2.96**$^*$ | **2.37**$^*$ | 4/5 | 2.89 |
| ECM organisation | **3.48**$^*$ | -1.06 | **2.62**$^*$ | **3.47**$^*$ | **3.56**$^*$ | 4/5 | 3.47 |
| KRAS signalling up | **3.21**$^*$ | **3.05**$^*$ | **2.56**$^*$ | **3.08**$^*$ | **1.91**$^*$ | 5/5 | 3.05 |
| TNF$\alpha$/NF-$\kappa$B | **2.00**$^*$ | **2.76**$^*$ | **2.14**$^*$ | **2.00**$^*$ | **1.77**$^*$ | 5/5 | 2.00 |
| **Between-method variability** | | | | | | | |
| EMT | **5.51**$^*$ | **3.39**$^*$ | **4.51**$^*$ | **5.36**$^*$ | **4.54**$^*$ | 5/5 | 4.54 |
| ECM organisation | **4.71**$^*$ | **2.67**$^*$ | **4.31**$^*$ | **5.35**$^*$ | **4.72**$^*$ | 5/5 | 4.71 |

**Mouse neural datasets**

| Programme | Allen MERFISH Pseudospot | Mouse Cortex Visium | Mouse Olfactory Bulb Rep12 | seqFISH+ Cortex/SVZ | STARmap Visual Cortex | $k/5$ | Median NES |
|---|---|---|---|---|---|---|---|
| **Mean PCC** | | | | | | | |
| Neuronal system | 1.29 | **4.65**$^*$ | **3.99**$^*$ | **2.15**$^*$ | **2.32**$^*$ | 4/5 | 2.32 |
| Synaptic signalling (GO BP) | **3.56**$^*$ | **5.95**$^*$ | **4.95**$^*$ | **3.29**$^*$ | **2.66**$^*$ | 5/5 | 3.56 |
| **Sparsity-adjusted mean PCC** | | | | | | | |
| Synaptic signalling (GO BP) | **3.27**$^*$ | **3.88**$^*$ | **3.48**$^*$ | **2.48**$^*$ | **2.31**$^*$ | 5/5 | 3.27 |
| **Between-method variability** | | | | | | | |
| Synaptic signalling (GO BP) | **2.82**$^*$ | **3.95**$^*$ | **3.33**$^*$ | **4.09**$^*$ | 1.09 | 4/5 | 3.33 |

**Interpretation.** Individual genes are not table rows: genes form the ranked input to GSEA, while each coloured cell is one programme-by-dataset result. For mean PCC, positive NES indicates enrichment among more reconstructable genes; for between-method variability, positive NES indicates greater instability across reconstruction methods. Human and mouse programmes were analysed separately using species-native collections.
